# Localisation and tissue tropism of the symbiont Microsporidia MB in the germ line and somatic tissues of Anopheles arabiensis

**DOI:** 10.1101/2023.03.06.531457

**Authors:** Edward E. Makhulu, Thomas O. Onchuru, Joseph Gichuhi, Fidel G. Otieno, Anne W. Wairimu, Joseph .N. Muthoni, Lizette Koekoemoer, Jeremy K. Herren

## Abstract

The *Anopheles* symbiont, *Microsporidia MB*, is maternally inherited and has a strong malaria transmission-blocking phenotype in *Anopheles arabiensis. Microsporidia MB* is also vertically transmitted, sexually transmitted and avirulent. These characteristics are expected to promote its spread through mosquito populations, enhancing the potential of *Microsporidia MB* as a candidate for the development of a symbiont-mediated malaria transmission blocking strategy. We found that the patterns of *Microsporidia MB* localisation over the development of *An. arabiensis* indicate accumulation in tissues linked to its transmission, specifically the male and female gonadal tissues. Transovarial vertical transmission of *Microsporidia MB* occurs in the female *An. arabiensis* ovary when *Microsporidia MB* becomes localised to the cytoplasm of the developing oocyte. In male *An. arabiensis, Microsporidia MB* is localised in the testis and vas deferens. Notably, a high intensity of *Microsporidia MB* can also be observed in the *An. arabiensis* adult but not larval gut. The levels of *Microsporidia MB* found in the female ovary are linked to the progression of oogenesis, increasing after blood feeding initiates the development of eggs. The levels of *Microsporiodia MB* in the male and female gonadal and gut tissue do not increase as mosquitoes age. Altogether, the high specificity of *Microsporidia MB* tissue localisation patterns and changes in infection prevalence and intensity suggest adaptation to maximise transmission and avirulence in *Anopheles arabiensis*.

**Importance:** *Microsporidia MB* is a symbiont with strong malaria transmission-blocking phenotype in *Anopheles arabiensis*. It spreads in mosquito populations through mother-to-offspring and sexual transmission. The ability of *Microsporidia MB* to block *Plasmodium* transmission together with its ability to spread within *Anopheles* populations and its avirulence to the host makes it a very attractive candidate for developing a key strategy to stop malaria transmissions. Here, we report the basis of *Microsporidia MB* transmission. We find that *Microsporidia MB* accumulates in *Anopheles arabiensis* tissues linked to its sexual and vertical transmission. Its prevalence and intensity in the tissues over the mosquito life cycle suggest adaptation to maximise transmission and avirulence in *Anopheles arabiensis*. These findings provide the foundation for understanding the factors that affect *Microsporidia MB* transmission efficiency. This will contribute to the establishment of strategies to maximize *Microsporidia MB* transmission for *Anopheles* mosquito population replacement and malaria transmission blocking.

## Introduction

The malaria disease burden remains a major impediment to good health and economic development in many regions of sub-Saharan Africa (SSA). In 2021, a total of 247 million cases were reported that resulted in 619,000 deaths, a strong indication that current control measures and their deployment levels are insufficient (1). Large-scale insecticide treated net (ITN) distribution campaigns have had a significant impact on reducing the number of malaria cases (2). However, many malaria vectors have now developed resistance to the insecticides used in ITNs (3) and they are increasingly biting outdoors, where nets offer no protection (4). In addition, many malaria control efforts were significantly disrupted by the COVID-19 pandemic (5) and the recent invasion of *An. stephensi* across Africa is leading to a significant rise in malaria transmission (6–8). Altogether, these factors threaten to reverse the gains achieved for malaria reduction and indicate an urgent need for new strategies to control *Anopheles* mosquito populations or their capacity to transmit *Plasmodium* parasites.

A novel and potentially transformative method of controlling vector-borne disease involves the use of transmission blocking symbionts. For dengue, a control strategy based on the transmission blocking symbiont *Wolbachia* has been highly effective and controlled field trials are currently implemented in over 13 countries (9). A similar strategy, based on a *Plasmodium* transmission-blocking symbiont in *Anopheles* mosquitoes could be transformative for controlling malaria. The microsporidian symbiont, *Microsporidia MB*, is naturally found in anopheline mosquito populations and has been shown to block *Plasmodium* development (10). In addition, *Microsporidia MB* is both vertically and sexually transmitted (11). In conjunction with the *Plasmodium* blocking, the high efficiency of vertical and sexual transmission of *Microsporidia MB*, which could facilitate the spread and maintenance of *Microsporidia MB* in *Anopheles* mosquito populations, has led to the suggestion that this symbiont could be deployed as a new tool for malaria control (12).

The success of a *Microsporidia MB-*based malaria control strategy will depend on efficient vertical and horizontal transmission of the symbiont, which would enable the spread and maintainance of *Microsporidia MB* in *Anopheles* vector populations. For symbionts that are vertically and sexually transmitted, avirulence towards the host can be expected (13). Vertically and sexually transmitted symbionts can be selected for their ability to enhance host fitness (14, 15). The tissue level localisation pattern and intensity of infection have been shown to play an important role in symbiont transmission and host fitness effects. Currently, little is known regarding *Microsporidia MB* localization patterns in infected mosquitoes. Here, we investigated the tissue-level localisation and changes of *Microsporidia MB* infection intensity across development of *An. arabiensis*. We found that *Microsporidia MB* was predominately found in the reproductive organs of *An. Arabiensis* males and females. Additionally, we observed *Microsporidia MB* inside developing oocytes in the *An. arabiensis* ovaries, indicating transovarial vertical transmission. Interestingly, the intensity of *Microsporidia MB* infection in the female ovaries increased after blood feeding. The prevalence and intensity of *Microsporidia MB* infection in the female ovaries was found to decrease as *An. arabiensis* mosquitoes aged. In male *An. arabiensis, Microsporidia MB* is found in the testis and vas deferens offering further confirmation of the basis of male to female sexual transmission as previously reported (11). *Microsporidia MB* is also in found in the *An. arabiensis* adult intestine at moderate prevalences. Notably, *Microsporidia MB* is always absent from the larval intestine but present in the larval body.

## Results

### The *An. arabiensis* male and female gonads are the primary site of *Microsporidia MB* infection in *An. arabiensis*

The prevalence and intensity of *Microsporidia MB* was investigated in the gonads, gut fat-body and carcass of seven-day-old adult *An. arabiensis* by qPCR (Figure 1). We show that *An. arabiensis* male and female gonads had the highest *Microsporidia MB* prevalence (Figure 1A) and the highest absolute intensity of *Microsporidia MB* infection in comparison to other tissues and the carcass (Figures 1B-1D). In female and male *An. arabiensis*, the gonads also had higher mean *Microsporidia MB* intensities relative to host gene copy number (relative *Microsporidia MB* intensity) than fat bodies and carcasses. These findings indicate that the primary site of *Microsporidia MB* infection in both male and female adult *An. arabiensis* was the gonadal tissue. This is in line with a study that investigated *Microsporidia MB* in only male *An. arabiensis* (11). A moderate prevalence of *Microsporidia MB* infection was observed in the *An. arabiensis* male gut (54%, Figure 1A). In addition, the mean relative *Microsporidia MB* intensity in the male gut was significantly higher than the fat body and carcass (Figure 1E), indicating that *Microsporidia MB* density can reach high levels in this tissue. In females, the relative intensity of *Microsporidia MB* in the gut was not significantly different from the relative intensity in the gonads (Figure 1C). Altogether, these findings suggest that the gut is likely to be the secondary site of infection. In both sexes, the prevalence and intensity of *Microsporidia MB* was lowest in the fat body (Figure 1A-E).

**Figure 1.**
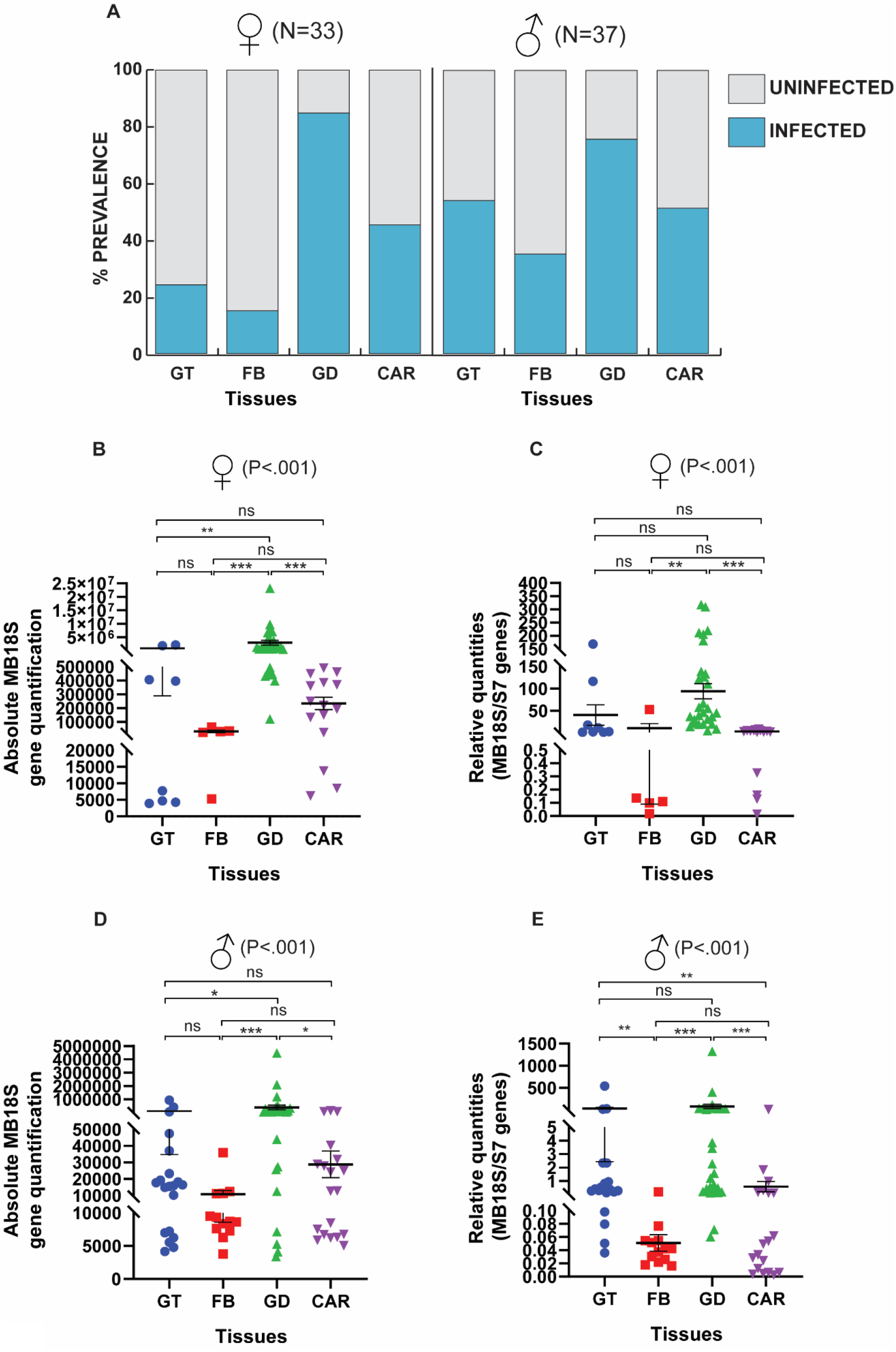
*Microsporidia MB* infection prevalence and intensity in adult *An. arabiensis* tissues. **(A)** The prevalence of *Microsporidia MB* is highest in the gonads of male and female *An. arabiensis* in comparison to other somatic tissues and the carcass. The absolute *Microsporidia MB* intensity was higher in female **(B)** (Dunn’s post hoc test, GT vs GD P =0.008, FB vs GD P<0.001, CAR vs GD P<0.001) and male **(C)** (Dunn’s post hoc test, GT vs GD P =0.019, FB vs GD P<0.001, CAR vs GD P =0.011) gonads relative to somatic tissues. **(D)** The relative intensity of Microsporidia MB in the female *An. arabiensis* midgut is not significantly different from the gonads. **(E)** The relative *Microsporidia MB* intensity determined by qPCR was significantly higher in the male gut in comparison to the fat body and carcass. (Abbreviations: **GT –** Gut, **FB –** Fat bodies, **GD –** Gonads, **CAR –** Carcass)

### *Microsporidia MB* cells are identified by microscopy in oocytes in female and testis and gut in male *An. arabiensis*

To better understand where *Microsporidia MB* is located inside the male and female *An. arabiensis* gonads and guts, we visualised these tissues using Fluorescent In-Situ Hybridization (FISH) and confocal microscopy. The visualisation of *Microsporidia MB* confirmed that male and females *An. arabiensis* gonads were the primary site of infection (Figure 2A-2B). In the *An. arabiensis* female gonad, *Microsporidia MB* is observed in all stages of egg chamber development as well as the germ-line stem cells (Figure 2A). As the oocytes enter vitellogenesis, higher intensities of *Microsporidia MB* are observed. In the male gonad, *Microsporidia MB* is observed in the testis and vas deferens, in close proximity to developing sperm cells (Figure 2B). In the *An. arabiensis* male gut, *Microsporidia MB* is occasionally observed at high infection levels in a small number of cells in the midgut and hindgut regions (Figure 2C).

**Figure 2.**
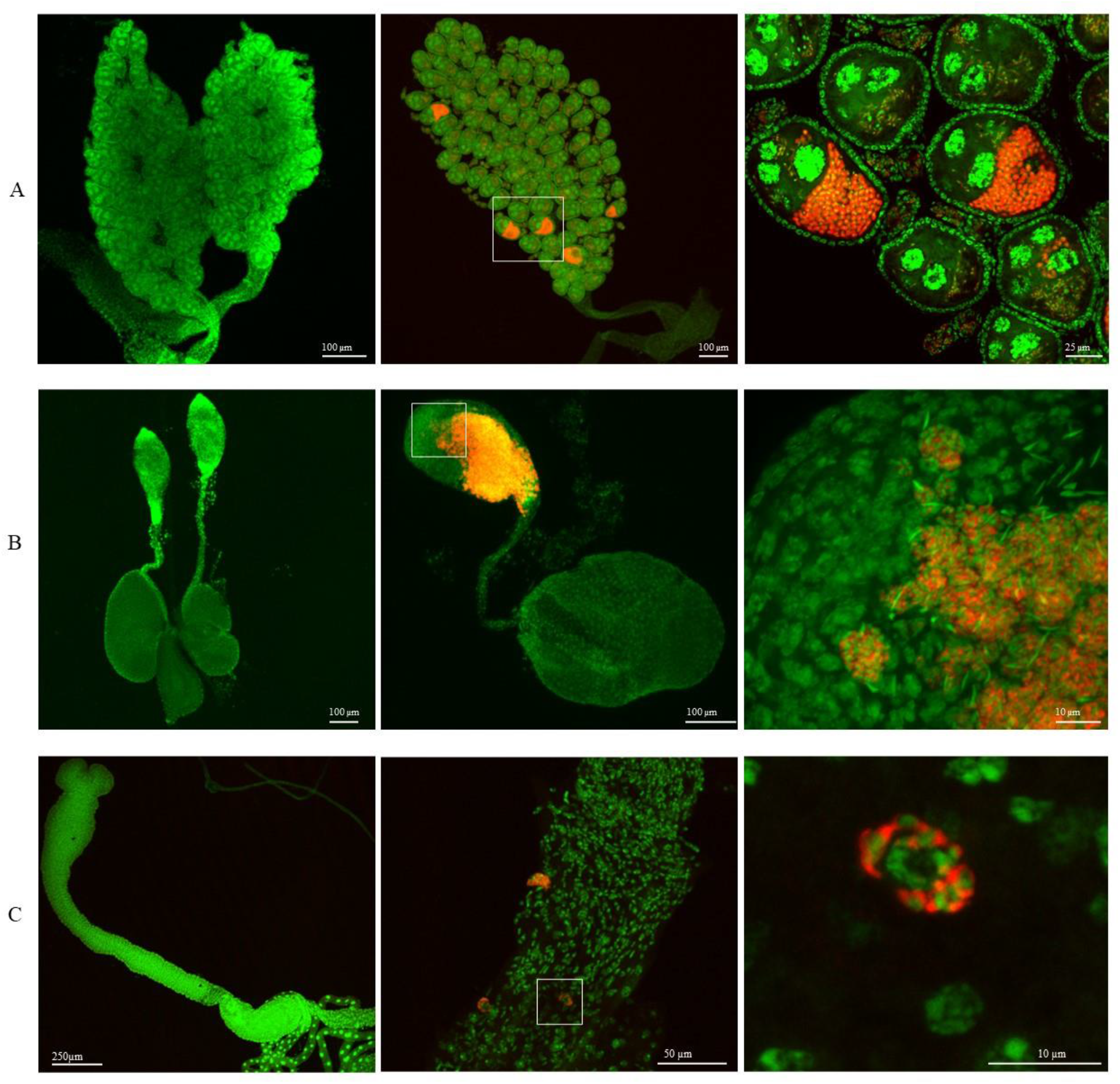
*Microsporidia MB* localisation to cells in the male and female gonads and male gut. *Microsporidia MB* infection is observed by FISH and confocal fluorescence microscopy in the gonads and gut of *An. arabiensis* mosquitoes. **(A)** *Microsporidia MB* in the female *An. arabiensis* gonads. A high density of *Microsporidia MB* is observed in vitellogenic oocytes. **(B)** *Microsporidia MB* in the male *An. arabiensis* gonad. A high density of *Microsporidia MB* is observed in parts of the testis. **(C)** *Microsporidia MB* is observed in the *An. arabiensis* male midgut cells.

### *Microsporidia MB* is not found in the larval gut and is higher in female adult mosquitoes

To better understand where *Microsporidia MB* is located during the larval stage of *An arabiensis*, L4 larval stages were dissected to separate the gut from the carcass, with carcasses containing all the remaining larval tissue, including the hemolymph. We observed that none of the gut samples were infected with *Microsporidia MB* (Figure 3A). However, larval carcasses were found to have a high prevalence of *Microsporidia MB* infection. The absolute intensity of *Microsporidia MB* infection in *An. arabiensis* larvae was significantly less than that observed in adult female but not male mosquitoes (Figure 3B).

**Figure 3.**
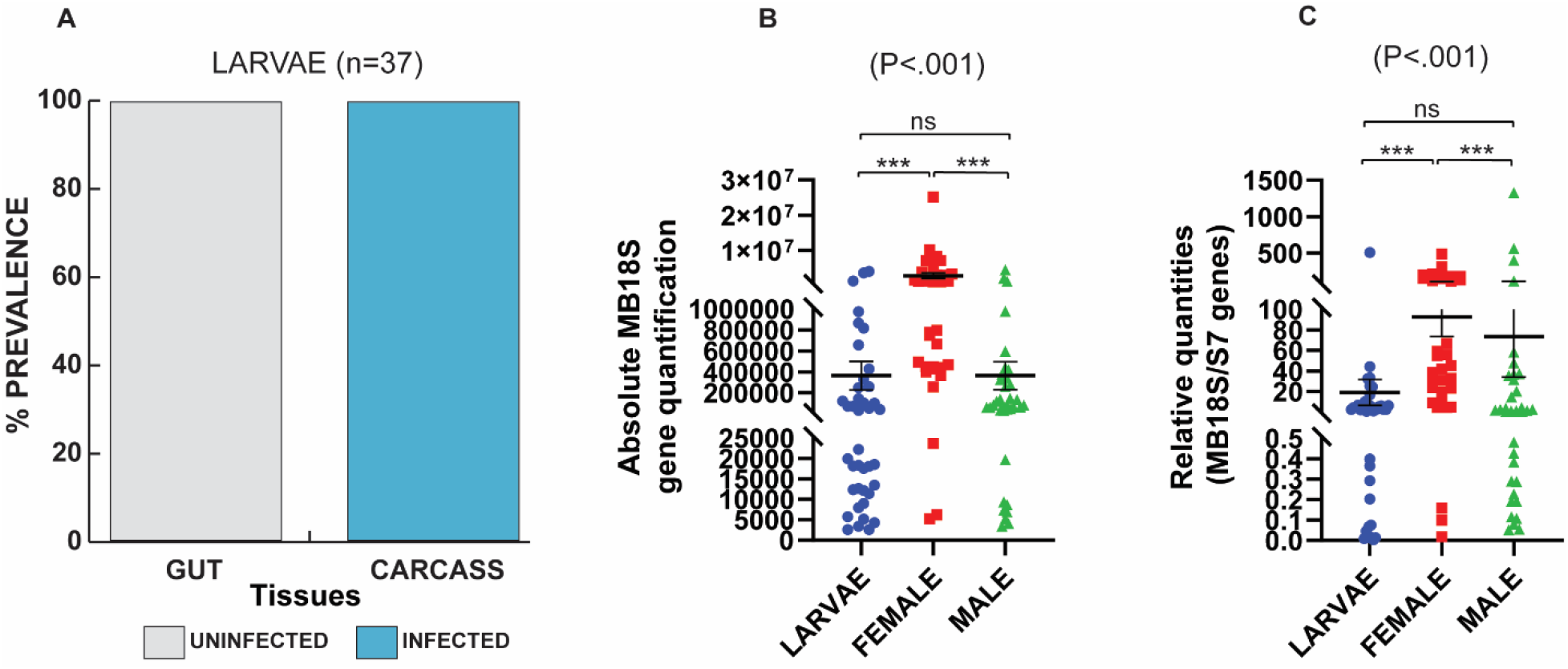
Localisation of *Microsporidia MB* in larval tissues and comparison of its density between larvae and adult mosquito stages. **(A)** *Microsporidia MB* is present in the carcass of dissected L4 larvae but not in the gut. *Microsporidia MB* infection intensity is only higher in *An. arabiensis* adult females relative to L4 larvae both in an absolute quantification (P<0.001) **(B)** and a relative quantification (P<0.001) **(C)** upon conducting a pairwise comparisons using Dunn’s test.

### Blood meal affects the intensity of *Microsporidia MB* in the ovaries

In *An. arabiensis*, oogenesis begins after female mosquitoes eclose but is arrested until females take a blood meal. To determine if the intensity of *Microsporidia MB* in *An. arabiensis* is affected by the onset of egg development and other physiological changes associated with taking a blood meal, we compared the prevalence and intensity of *Microsporidia MB* infection in the guts and gonads of *An. arabiensis* females that had fed on a blood meal with those that had not fed on a blood meal. In the *An. arabiensis* female gut, the prevalence of *Microsporidia MB* was slightly lower in blood fed mosquitoes relative to non-blood fed mosquitoes (Figure 4A). While there was a significantly higher relative intensity of *Microsporidia MB* in female *An. arabiensis* guts that had been blood fed, the absolute levels of *Microsporidia MB* were not significantly different between the two groups (Figure 4C and 4E). This discrepancy is likely the result of the turnover (and shedding) of gut epithelial cells that is known to occur after blood feeding, which could decrease the amount of host DNA in this tissue (16,17). The prevalence of *Microsporidia MB* infection was high in ovaries regardless of the blood feeding status (Figure 4B). However, we observed significantly higher relative and absolute intensities of *Microsporidia MB* in gonads of blood fed *An. arabiensis* (Figure 4D and 4F). These observations suggests that the amount and density of *Microsporidia MB* in the female gonad increases as the gonotrophic cycle progresses and egg development is initiated.

**Figure 4.**
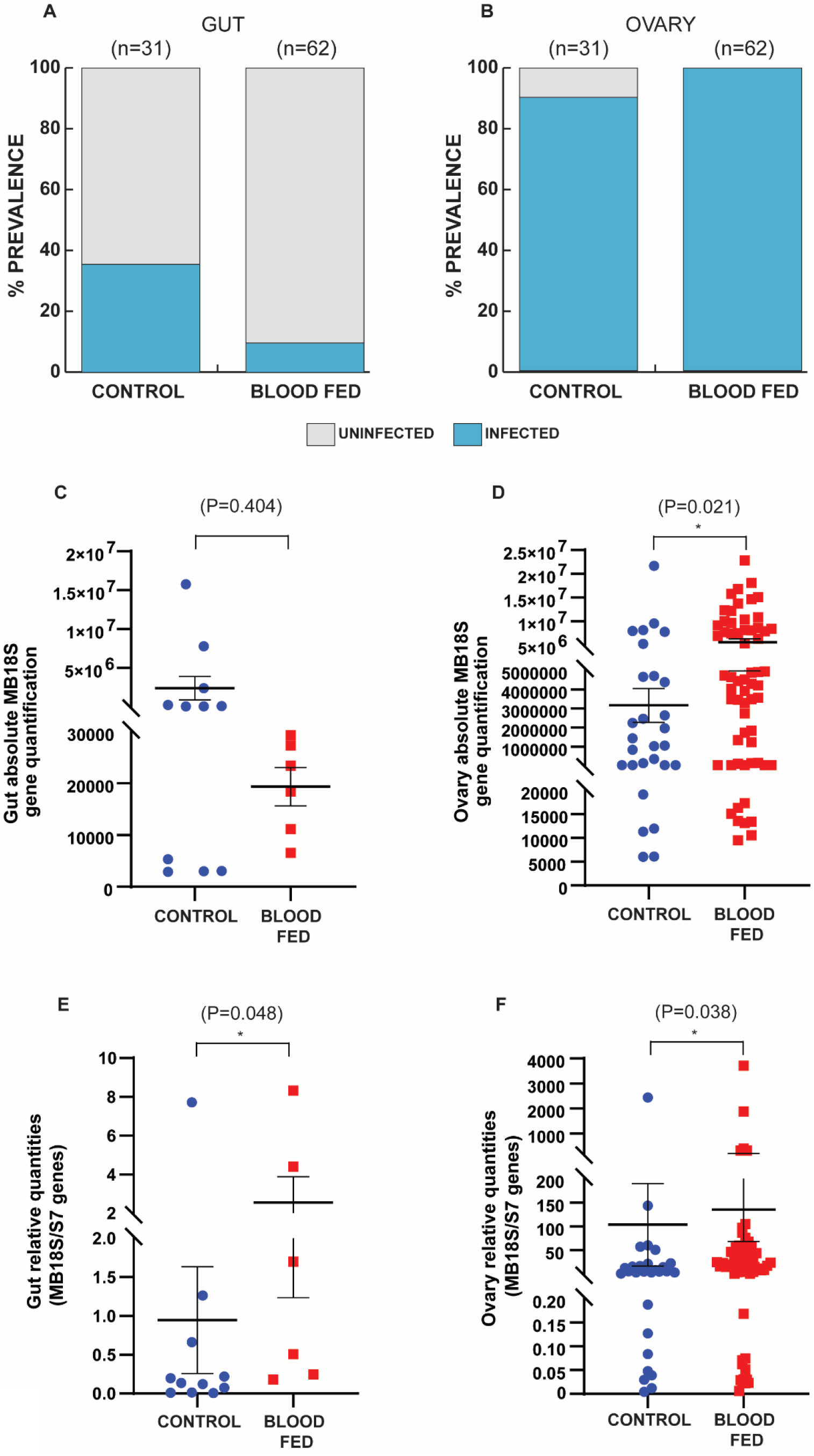
*Microsporidia MB* prevalence and intensity in female *An. arabiensis* gonads is affected by blood feeding. **(A)** The prevalence of *Microsporidia MB* infection is slightly lower in the guts of blood fed *An. arabiensis* females. **(B)** The prevalence of *Microsporidia MB* infection is high in blood fed and non-blood fed *An. arabiensis* female gonads. **(C)** The absolute intensity of *Microsporidia MB* infection is not significantly different between the guts of blood fed and non-blood fed *An. arabiensis* females (two-tailed Mann–Whitney test, P = 0.404, error bars reflect SEM). **(D)** The absolute intensity of *Microsporidia MB* infection is significantly higher in the gonads of blood fed *An. arabiensis* females (two-tailed Mann–Whitney test, P = 0.021, error bars reflect SEM). **(E)** The relative intensity of *Microsporidia MB* infection is significantly higher in the guts of blood fed *An. arabiensis* females (two-tailed Mann–Whitney test, P = 0.048, error bars reflect SEM). **(F)** The relative intensity of *Microsporidia MB* infection is higher in the gonads of blood fed *An. arabiensis* females (two-tailed Mann–Whitney test, P = 0.038, error bars reflect SEM).

**Figure 5.**
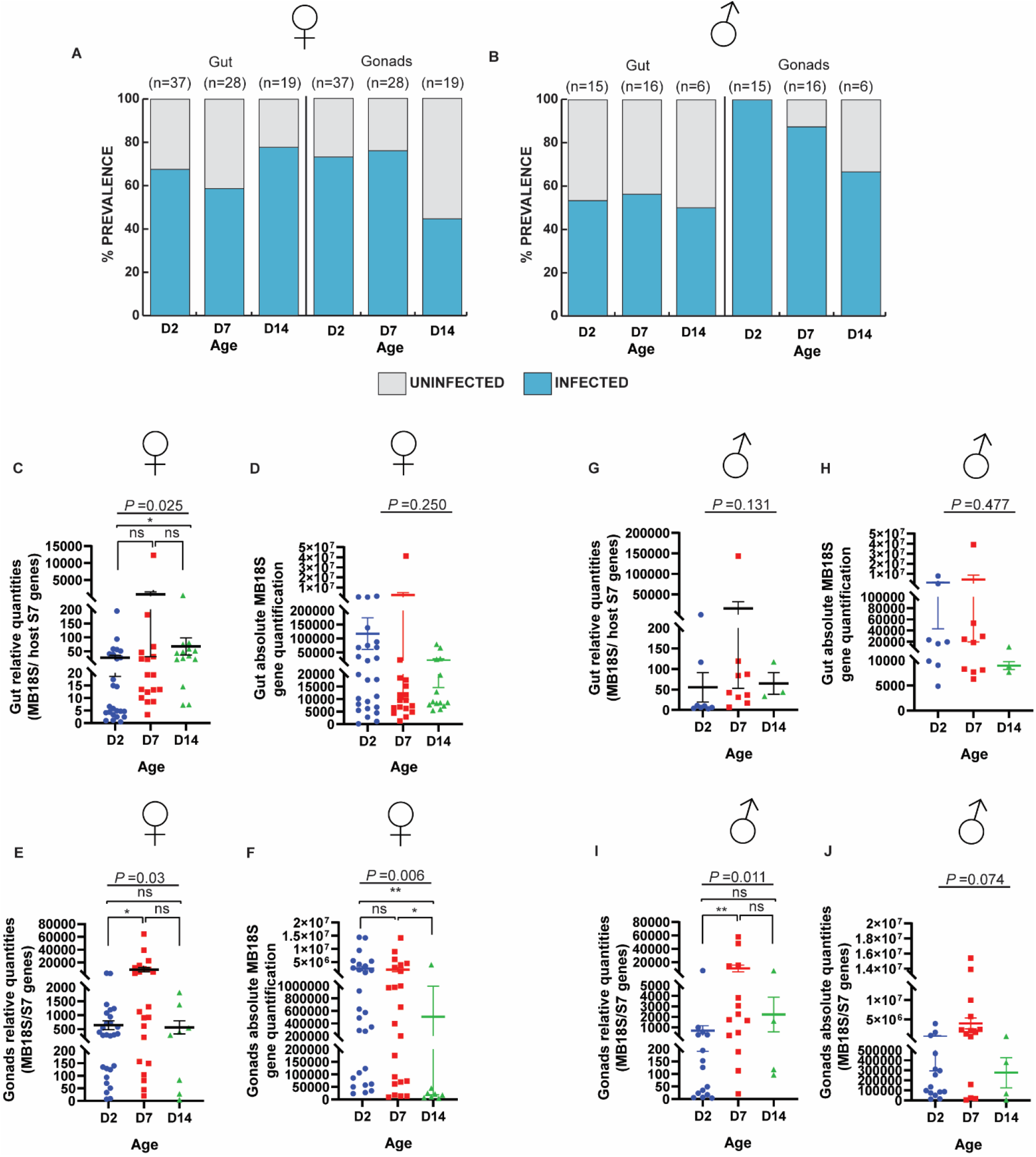
The intensity of *Microsporidia MB* in the *An. arabiensis* gonad and gut is affected by age. **(A)** The prevalence of *Microsporidia MB* in the guts of female *An. arabiensis* was slightly higher at 14 days relative to seven and two days. A decrease in the prevalence of *Microsporidia MB* is observed in the gonads of 14-day old female *An. arabiensis*. **(B)** The prevalence of *Microsporidia MB* in the guts of male *An. arabiensis* did not change as mosquitoes were aged. A decrease in the prevalence of *Microsporidia MB* is observed in the gonads of 14-day old male *An. arabiensis*. **(C-D)** An increase in the relative intensity (Dunn’s post hoc test, P =0.022), but not absolute intensity, of *Microsporidia MB* is observed in the guts of female *An. arabiensis* between 2- and 14 days. **(E-F)** The relative intensity (Dunn’s post hoc test, P =0.046), but not absolute intensity, of *Microsporidia MB* in female *An. arabiensis* gonads increased between day two and seven. The absolute intensity of *Microsporidia MB* was lower in the gonads of 14-day old females relative to (Dunn’s post hoc test, P =0.005) and seven (Dunn’s post hoc test, P =0.022) females. **(G-H)** The intensity of *Microsporidia MB* in the guts of male *An. arabiensis* does not change over aging. **(I-J)** The relative intensity of *Microsporidia MB* in the *An. arabiensis* male gonad increased between two and seven days of aging (Dunn’s post hoc test, P =0.008). The absolute intensity of *Microsporidia MB* in the male gonad was not significantly different across all ages.

### Age affects the intensity of *Microsporidia MB* density in *An. arabiensis* gonads and guts

To investigate the effect of aging on the prevalence and intensity of *Microsporidia MB* in *An. arabiensis* gonadal and gut tissue, male and female *An. arabiensis* were aged for 2, 7 and 14 days prior to dissection of tissues and quantification of *Microsporidia MB* by qPCR. The prevalence of *Microsporidia MB* in the guts of female *An. arabiensis* was slightly higher in the 14-day old mosquitoes relative to 2- and 7-day old mosquitoes (Figure 4A). The prevalence of *Microsporidia MB* in the guts of male *An. arabiensis* did not change as mosquitoes were aged (Figure 4B). In both males and females, there was a general trend towards a decrease in the prevalence of *Microsporidia MB* in gonads as mosquitoes were aged (Figure 4A-4B). There was a slight increase in the relative intensity, but not absolute intensity of *Microsporidia MB* in the guts of female *An. arabiensis* between 2- and 14 days (Figure 4C-4D). The intensity of *Microsporidia MB* in the guts of male *An. arabiensis* was apparently not affected by aging (Figure 4G-4H). The relative intensity of *Microsporidia MB* in female *An. arabiensis* gonads increased between day 2 and 7, however, the absolute levels remained constant (Figure 4E-4F). This suggests that the increase in relative *Microsporidia MB* intensity is a consequence of host DNA content decreasing. Notably, the absolute, but not relative, intensity of *Microsporidia MB* was lower in the gonads of 14 day old females relative to 2 and 7 day old females (Figure 4E-4F). The relative intensity of *Microsporidia MB* in the *An. arabiensis* male gonad increased between 2 and 7 days of aging (Figure 4I). The absolute intensity of *Microsporidia MB* in the male gonad was not significantly different across all ages (Figure 4J).

## Discussion

The results demonstrate that the gonads of male and female *An. arabiensis* are the primary sites of *Microsporidia MB* infection. This finding indicates that maternal and sexual transmission are likely to be the most important *Microsporidia MB* infection routes. Both maternal and sexual transmission require that *An. arabiensis* hosts do not incur excessive fitness costs when infected with *Microsporidia MB* (13). In line with this, *Microsporidia MB* does not have a major effect on the fitness of larvae or adult female mosquitoes (10). It remains unresolved how *Microsporidia MB* can reach very high intensities, up to 2.5×10^7^ copies of the *Microsporida MB* 18S gene per mosquito, without negatively impacting host fitness.

Other insect endosymbionts, including *Wolbachia*, have intensity-dependent effects on host fitness (18). It is therefore possible that very high intensity *Microsporidia MB* infections do have a fitness cost, but this would need further investigation. The link between *Wolbachia* intensity and host fitness is complex and, in many cases, fitness is not affected by high endosymbiont intensities (19). It is likely that the pattern of endosymbiont localisation in host tissues has important fitness consequences for endosymbionts such as *Wolbachia* and *Microsporidia MB*. A strain of *Wolbachia, w*MelPop, that over proliferates and decreases host fitness has been found at high intensities in the central nervous system and muscles of its insect hosts (20, 21). It is possible that endosymbionts mitigate effects on host fitness by limiting most of the infection and proliferation to certain tissues.

Our findings indicate that *Microsporidia MB* is primarily found in gonadal tissue and this is suggestive adaptation to minimise host fitness costs without compromising transmission. Fluorescence confocal microscopy revealed that *Microsporidia MB* was present in oocytes and nurse cells across all stages of oogenesis. These findings clearly demonstrate that *Microsporidia MB* maternal transmission is transovarial, which explains the need for higher *Microsporidia MB* intensities to be maintained in the female gonads.

In the male germline, *Microsporidia MB* is primarily localised to the testis and vas deferens. The testis is the site of sperm production, comprising of a proximal end with stem cell divisions, spermatocysts in different stages of development and a distal spermatozoa reservoir (22). In *An. arabiensis*, we observed *Microsporidia MB* cells as clusters in the spermatocyst and spermatozoa reservoir regions of the testis and more rarely in the stem cell region. These findings suggest that *Microsporidia MB* is packaged with spermatozoa at their early development stages.

We observed very low levels of *Microsporidia MB* in the *An. arabiensis* adult fat body. Pathogenic Microsporidians are known to infect and proliferate in the insect fat body (22). It is likely that as the primary energy storage tissue, fat bodies provide the nutrients required for this proliferation. The low levels of *Microsporidia MB* in the adult fat body of *An. arabiensis* could be part of the explanation for its avirulence in *An. arabiensis*. The moderate levels of *Microsporidia MB* in the carcass samples, which contained all the tissues apart from gut, gonads and fat body, suggest that we cannot exclude there being another *Microsporidia MB* infection reservoir in adult *An. arabiensis*.

In the adult *An. arabiensis* gut, there were a small proportion of samples which had very high intensities of *Microsporidia MB*. The significance of these high intensity gut infections is yet to be resolved. Fluorescence confocal microscopy revealed that *Microsporidia MB* was found at very high densities in a small subset of cells in the adult male mid and hindgut. The shape and positioning of these cells suggests that they could be intestinal stem cells (17). It is notable that in contrast to adults, *Microsporidia MB* was never observed in the larval gut. A possible explanation for this is that presence in the larval gut could have prohibitive fitness cost since larvae feed continuously. In addition, the total intensity of *Microsporidia MB* is lower in larvae relative to adult females.

We also investigated how localisation and *Microsporidia MB* intensity changes over the life cycle of *An. arabiensis*. In *An. arabiensis*, oogenesis starts when mosquitoes elcose, but is then paused until females take a blood meal (24). Once females take a blood meal, several physiological changes occur that enable the blood to be digested an ultimately to meet nutritional demands of egg production (25). Since there are major changes occurring in the gut and gonads of females post blood feeding, we investigated the changes in the levels of *Microsporidia MB* in these tissues. We observed that the levels of *Microsporidia MB* in the gut did not change in response to blood feeding; the relative intensity of *Microsporidia MB* did increase in the gut after blood feeding but the absolute intensity appeared to be constant, which suggests a decrease in host nuclear gene copy number. This decrease is likely to be linked to the process of gut epithelium renewal (17). In contrast, we observed a marked increase in the intensity of *Microsporidia MB* in the female gonad after blood feeding. Notably, *Microsporidia MB* is observed at high intensities in the later (vitellogenic) stages of oogenic development, which develop after the blood meal induced initiation of oogenesis (24). Therefore, the proliferation of *Microsporidia MB* in vitellogenic oocytes is a possible explanation for the increase in intensity of *Microsporidia MB* after blood feeding.

We did not observe *Microsporidia MB* levels increasing as male or female guts and gonads as *An. arabiensis* mosquitoes are aged. This pattern of growth is consistent with symbionts that regulate proliferation to mitigate fitness costs (26). While there was an increase in the relative intensity of *Microsporida MB* in male and female gonads between day 2 and day 7, this is likely to be the consequence of a decrease in host nuclear gene copy number. Notably, there was a significant decrease in the prevalence and absolute intensity of *Microsporidia MB* in female gonads between 7 and 14 days of aging. This may be indicating a loss of *Microsporidia MB* from germline cells after numerous rounds of division and an inability for these cell to be re-infected. However, the aged *An. arabiensis* female mosquitoes had not been blood fed and therefore had not completed any gonotrophic cycles which could be expected to lower the rate of stem cell division (27).

Overall, our findings suggest that *Microsporidia MB* localisation patterns may serve to maximise transmission success while minimising negative effects on host fitness. *Microsporidia MB* has proliferation and localisation patterns that are consistent with a co-evolved symbiont that exploits sexual and vertical transmission routes. A better understanding of changes in localisation and symbiont intensity across the *An. arabiensis* lifecycle will contribute to improving *Microsporidia MB* infected mosquito rearing and aid in the formulation of strategies to disseminate *Microsporidia MB*.

## Materials and methods

### Mosquito sample collection

Wild-caught gravid adult female *Anopheles gambiae s*.*l*., were collected using mouth aspiration. Collected mosquitoes were identified morphologically to the species level and sub-species identification of *Anopheles gambiae s*.*l*., was carried out by PCR (28). Sampling was conducted from one site in Kenya, the Ahero irrigation scheme (– 34.9190W, –0.1661N). The mosquitoes used in this study were collected between September 2021 and October 2021. Sampled mosquitoes were transported to the *icipe* - Duduville campus in Nairobi and maintained on 6% glucose and water.

### Mosquito processing and development of isofemale lines

Wild collected mosquitoes were maintained in an insectary at 27 ± 2.5°C, humidity 60– 80% and 12-h day and 12-h night cycles and induced to oviposit in individual microcentrifuge tubes containing a wet 1 cm × 1 cm Whatman filter paper. Eggs from each female were counted under a compound microscope using a paint brush and then dispensed into water troughs for larval development at 30.5°C and 30–40% humidity. Tetramin^™^ baby fish food was used to feed developing larvae. Upon laying eggs, the G^0^ females were screened for presence of *Microsporidia MB* by PCR. The larval offspring of *Microsporidia MB* positive field-caught female mosquitoes were transferred into larval rearing troughs for further development. Upon reaching the L4 larval stage, representative samples from each positive isofemale line were screened to determine the *Microsporidia MB* infection status of the line. Were mosquitos were maintained as adults, this was done in an insectary at 30°C, with a relative humidity of 75% and 12-h day and 12-h night cycles and a feeder with 6% glucose solution.

### Larval and adult dissection

*An. arabiensis* G1 larvae and adult mosquitoes were dissected using forceps under the Zeiss Stemi 2000-C stereomicroscope to obtain G1 L4 larval and adult *Anopheles* tissues. During the dissection of the L4 larvae, alive samples were placed on a drop of 1X Phosphate Buffered Saline (1X PBS). The larvae were restrained at the junction between the head and thorax with a pair of forceps. A dissecting needle pin was then used to probe the intersection between the second and last abdomen segments. The siphon and saddle of the larvae were held and pulled gently to obtain the gut of the larvae using a new pair of forceps. The carcass and the gut were placed in separate 1.5 micro-centrifuge tubes containing 20µl of 1X PBS and 0.5mm zirconium beads for DNA extractions. For adults, G1 mosquito samples were first anaesthetised for a few minutes until immobilised by aspirating them in 1.5 micro-centrifuge tubes and placing them in ice. Upon immobilisation, the mosquitoes were placed in a drop of 1X PBS on a glass slide. The junction between the last and second-last abdomen was probed using a dissecting needle. The last segment was gently pulled to release the gonads and the gut. The gut and gonads were separated and placed in separate 1.5 micro-centrifuge tubes containing 20µl of 1X PBS and 0.5mm zirconium beads ready for nucleic acid extractions. Similarly, the fat bodies and carcass (all remaining tissue) and were also separated and placed in separate 1.5 micro-centrifuge tubes as described above. Forceps were sterilised after every dissection to prevent contamination.

### Mosquito blood feeding

Two to three-day-old *Microsporidia MB* positive and negative *An. arabiensis* mosquitoes maintained at a temperature of 30°C, and a relative humidity of 75 % were starved for 2 hours without water or glucose in preparation for blood feeding. Membrane feeding was conducted according to (29). Breifly, bovine blood was transferred into a Hemotek® feeding apparatus (Hemotek, UK) with a Parafilm-A membrane set at. The Hemotek feeding apparatus was placed on top of a cage of starved *An. arabiensis* mosquitoes and feeding was allowed to take place for 1hr. Fully engorged *An. arabiensis* were transferred to new cages, maintained for five days post-infection, and later dissected to obtain the gut and ovaries which were screened for *Microsporidia MB* presence and levels.

### Flourescent *in-situ* hybridisation (FISH) localisation

Dissected tissues were fixed overnight in 4% Paraformaldehyde (PFA) at 4 °C. After fixation, the samples were rehydrated in 50% ethanol for 30 minutes and transferred to 1X PBS for another 30 minutes. FISH was then conducted to localize *Microsporia MB* within the tissues. Hybridization was done by incubating the tissues in 100μl of hybridization buffer overnight at 50°C. The FISH hybridization mix contained hybridization buffer (dH2O, 5M NaCl, 1M Tris/HCl [pH=8], and 10% SDS), *Microsporidia MB* specific CY5 probe (10), 0.5μM final concentration, and SYTOX Green general DNA staining. After staining, the samples were washed twice with 100μl of prewarmed wash buffer (dH2O, 5M NaCl, 1M Tris/HCl [pH=8], 0.5M EDTA, and 10% SDS). The tissues were then placed on a slide and visualized immediately using a Leica SP5 confocal microscope (Leica Microsystems, USA). The images were analyzed with the ImageJ 1.50i software package (30).

### DNA extraction and molecular detection and quantification of *Microsporidia MB*

DNA was extracted using the ammonium acetate protein precipitation method (10). Extracted DNA samples were screened to determine their *Microsporidia MB* infection status using the *Microsporidia MB* specific primers (MB18SF: CGCCGG CCGTGAAAAATTTA and MB18SR: CCTTGGACGTG GGAGCTATC) targeting the *Microsporidia MB* 18S rRNA region (10). The PCR reaction used in detection comprised a 10 µl reaction consisting of 2 µl HOTFirepol Blend Master mix Ready-To-Load (Solis Biodyne, Estonia), 0.5 µl of 5 pmol/µl forward and reverse primers, 2 µl of the DNA template, and 5 µl nuclease-free PCR water was undertaken. The thermocycling conditions employed were initial denaturation at 95°C for 15 min, followed by 35 cycles of denaturation at 95°C for 1 min, annealing at 62°C for 90 s, and extension at 72°C for a further 60 s. Final elongation was done at 72°C for 5 min. Samples positive for *Microsporidia MB* were subjected to relative and absolute qPCR analysis to quantify infection levels (10, 31). The qPCR analysis involved the MB18SF/ MB18SR primers, normalised with the *Anopheles* ribosomal S7 gene (primers, S7F: TCCTGGAGCTGGAGATGAAC and S7R: GACGGGTCTGTACCTTCTGG) as the reference host gene. The 10µl PCR reaction consisted of 2µl HOT FIREPol® EvaGreen® HRM no ROX Mix Solis qPCR Master mix (Solis Biodyne, Estonia), 0.5 µl of 5 pmol/µl forward and reverse primers, 2 µl of the DNA template from *Microsporidia MB* positive samples, and 5 µl nuclease-free PCR. The thermocycling conditions employed were an initial denaturation at 95°C for 15 min, followed by 35 cycles of denaturation at 95°C for 1 min, annealing at 62°C for 90 s, and extension at 72°C for a further 60 s. Finally, melting curves were generated by melting analysis with temperature ranges from 65°C to 95°C. The PCR and qPCR was carried out on a MIC qPCR cycler (BioMolecular Systems, Australia). Each sample was confirmed to have the characteristic melt curve associated with the *Microsporidia MB* MB18SF/ MB18SR primers.

### Data analysis

All the data was tested for nomarlity using the Shapiro-Wilk test. For non-normal unpaired data, a two tailed Mann–Whitney U test was used to determine the difference between the groups. In cases with more than two data groups, we used Kruskal-Wallis H test to determine the significance of differences between the groups. If significant, we carried out a Dunn’s post hoc test. All statistical analyses were preformed using Graphpad Prizm version 6.0c software or R (version 8.0.2). P-values of *p < 0.05, **p < 0.01, ***p < 0.001 and ****p < 0.0001 were deemed to be statistically significant. Figures were created and/or edited using Adobe Illustrator (version 23.0.1).

## Acknowledgement

The study was supported by Open philanthropy (grant no.). The funders had no role in study design, data collection, and analysis, decision to publish, or preparation of the manuscript. We also thank the support provided by the *icipe* insectary team (Milcah Gitau, Jeniffer Thiong’o, Peris Wambui, David Alila, and Charles Amara), field work team (Gerald Ronoh and Robinson Kisero) and the project administrator Faith Kyengo.

## Data accessibility

The following data will be submitted to the Dryad repository: *Microsporidia MB* localization and prevalence in adult tissues, prevalence and density in larvae vs adults, effect of bloodfeeding on prevalence and density of *Microsporidia MB*, density of *Microsporidia MB* in the *An. arabiensis* gonad and gut across different ages.

## Author contributions

EEM – Conceptualization, data curation, validation, visualization, formal analysis, investigation, methodology, writing – original draft, writing – review & editing.

TOO – Data curation, validation, visualization, formal analysis, investigation, methodology, supervision, writing – original draft, writing – review & editing.

JG, FGO, AWW, JNM – Investigation, writing – review & editing. LK – Conceptualization, supervision, writing – review & editing.

JKH – Conceptualization, data curation, formal analysis, funding acquisition, methodology, supervision, validation, visualization, writing – original draft, writing – review & editing.

